## Supplementary Material for "Differential quantification of alternative splicing events on spliced pangenome graphs"

*Simone Ciccolella, Davide Cozzi, Gianluca Della Vedova, Stephen Njuguna Kuria, Paola Bonizzoni, and Luca Denti*

### S1 Spliced pangenome construction and alignment

`pantas` requires as input an annotated spliced pangenomes (in GFA format) and RNA-Seq alignments to this structure (in GAF format). However, since these files are not always readily available, we implemented a custom procedure based on the `vg` toolkit [GSN$^+$18] to prepare these inputs. We decided to use `vg` since it is the only framework that currently support spliced alignments to a pangenome. In order to have full control of each step of its construction and be able to annotate the graph with additional information needed for AS events discovery (hence making an *annotated spliced pangenome*), we opted for manually constructing and indexing the graph. Indeed, although `vg` allows a user to automatically build an index via the `autoindex` utility, it does not allow to easily retain information on transcripts and splice junctions, that are needed for correct AS event detection. As suggested by `vg` authors, our procedure starts by chunking the input files (reference genome, gene annotation, and variations) by chromosomes and builds one graph per chromosome. It uses `vg construct` to build a pangenome from the reference chromosome and the set of variations and then it augments the graph with transcripts and splice junctions information using the `vg rna` utility. The resulting spliced pangenome contains new edges representing the annotated splice junctions but does not contain the haplotype-specific transcripts yet. Indeed, as suggested by `vg` authors, to maintain the construction as efficient as possible, it is more convenient to first build the spliced pangenome, extract the full haplotypes from the graph (using `vg gbwt`), and finally rerun the `vg rna` utility to project the reference transcripts against the haplotypes, thus obtaining the set of haplotype-aware transcripts. This is the methodology implemented in `pantas`. Once a spliced pangenome with haplotype-aware transcripts is available, `pantas` proceeds to simplify it to remove complex regions and improve the feasibility of the indexing step. `pantas` provides three different simplification steps. The default simplification consists in pruning only the intergenic and intronic regions of the graph. This is done by including all haplotype-aware trascripts and then using the `vg prune` utility asking to restore all paths (i.e., all transcripts) in the graph (setting the optional argument `--restore-paths`). A more aggressive simplification consists in forcing the inclusion of only the reference transcripts. Similarly to the default simplification, this is done by including the reference transcripts in the graph and then running the `vg prune` utility. With this second alternative, exonic portions of the genes could be simplified if too complex. The last simplification consists in removing all intergenic regions and thus retaining only the genic loci. The final graph is a set of connected components, each one representing a gene locus. We notice that a gene locus can potentially include multiple genes in the case of overlapping genes. The choice of which simplification to perform is use case dependent and needs to be decided by the user. The default simplification should provide the most accurate results (since all haplotype-aware transcripts are retained) but indexing the resulting graph will result very computationally expensive. On the other hand, the more aggressive simplification produces a graph that is easier to index, at the expense of potentially lower precision in calling haplotype-aware events (since some haplotype-aware transcripts may be removed and then not annotated). Finally, the third simplification provides the smallest graph while maintaining full information on haplotype-aware transcripts but it is especially useful when the user is interest in analyzing a small panel of genes (as it is the case in our experimental evaluation on real data).

After simplifying the graph, our procedure proceeds to annotate the vertices and the edges of the graph with additional information. Namely, it tags each vertex belonging to an exon with the name of the transcript it belongs to and the exon number (along that transcript) whereas each edge representing a splice junction with the number of the two linked exons and the transcript. The annotation iterates over the haplotype-aware transcripts retained in the graph and tag its vertices. Every time an edge is not present in the corresponding haplotype, it is by definition a splice junction and is tagged as such. We note that vertices and junctions can be shared by more transcripts and therefore annotated with multiple tags (i.e., a set). This information is a valuable information needed for AS events inference from graph alignment. These chromosome-level annotated spliced pangenome graphs are then merged together (without modifying the vertex id space) and the GCSA2 index of the resulting graph is built using the `vg index` utility. We note that spliced pangenome construction and indexing is a one-time expense and the index can be reused for any number of RNA-Seq datasets.

Once an annotated spliced pangenome index is available, the input RNA-Seq dataset can be aligned to it using the `mpmap` aligner [SEN+23]. This aligner was specifically introduced to align RNA-Seq to spliced pangenomes and to this aim it can align over novel splice junctions, that are edges not present in the input graph. This feature is essential for inferring novel AS events. The pipeline to prepare the input for `pantas` is provided alongside the main `pantas` codebase at `github.com/algolab/pantas`.

### S2 Details on the simulation settings

To evaluate `pantas` on simulated data, we considered the Drosophila Melanogaster reference and gene annotation (FlyBase [TGS+19], r6.51) and the Drosophila Genetic Reference Panel (v2) [MRS+12]. To avoid unnecessary complexity from our exploratory analysis on simulated data, we removed all overlapping genes from the gene annotation and we considered only SNPs (Single Nucleotide Polymorphisms) falling in the resulting set of genes with minimum count and frequency for non-reference allele equal to 1 and 0.01, respectively. The resulting reference panel consists of 2 311 679 SNPs and 205 inbred lines (samples). After randomly selecting one sample from the reference panel, we created its genomic sequence using `bcftools consensus` [DBL+21] and then we simulated a RNA-Seq dataset (2 conditions, 1 replicate) using `asimulator` [MTF+21]. We generated 25 000 000 read pairs and we enabled the simulation of exon skipping, alternative acceptor, alternative donor, and intron retention events (with a maximum of 1 event per gene). `asimulator` works by selecting a transcript as *template* transcript and, starting from this, it introduces AS events by altering the template and creating additional *alternate* transcripts. In doing so, it creates a new annotation containing all the generated transcripts. In our analysis, we considered this annotation as input for the tested tools and not the original gene annotation since `asimulator` may merge different transcripts in a single template transcript, thus creating chimeric transcripts not present in the input annotation. Starting from the `asimulator` annotation, which contains all transcripts used to simulate reads (template transcript and alternate transcripts), we also created a reduced annotation containing the template transcript only. This reduced annotation is used to simulate a poor annotation and test the accuracy of tools in calling novel AS events. Based on this data generation, we evaluated the accuracy of `pantas`, `rMATS` [SPL+14], `SUPPA2` [TEH+18], and `whippet` [SWWB+18]) in terms of Precision, Recall, and F1-Measure computed by comparing the events reported by `asimulator` with the events reported by the tools. In the annotated setting, we considered an event to be a *True Positive* if the reported splice junctions are correct with respect to the junctions reported by the simulator. In the novel setting, instead, due to the higher complexity of correctly detecting the novel splice junction, we considered an event to be a *True Positive* if at least one splice junction involved in the event is the correct one. Given the noise present in the truth created by `asimulator` (i.e., events barely supported and events with very low differential expression), we retained only those events with $|\Delta\psi| \geq 0.05$ and we created several truth sets filtering out all events supported by less than $\omega$ reads, with $\omega \in \{1, 3, 5, 10, 20\}$. For instance, with $\omega = 3$, we filtered out all events reported by `asimulator` supported by less than ($<$) 3 reads. Therefore, a bigger $\omega$ value results in a smaller sets of highly supported events, as also reported in Suppl.Table S1.

### S3 RT-PCR truthset filtering

The original RT-PCR truthset provided by [TEH+18] consists of 83 events on 82 genes. However, we note that the original RT-PCR validated events are reported on the `hg19` reference genome. We then used the `liftover` utility to convert their coordinates to the newer `hg38`. However, for three of the 83 events, the remapped coordinates disagreed with the gene annotation (due to a change in start/end position of the skipped exons). We then filtered these three events out and focused our analysis on the resulting set of 80 events. From this set of RT-PCR events, we also removed all events reported with low differential change between the two conditions and we considered all events with $|\Delta\psi| \geq 0.05$. This filter removed 3 additional events from the truth set. The resulting truthset consists in 77 RT-PCR validated events.

### Supplementary Figures and Tables

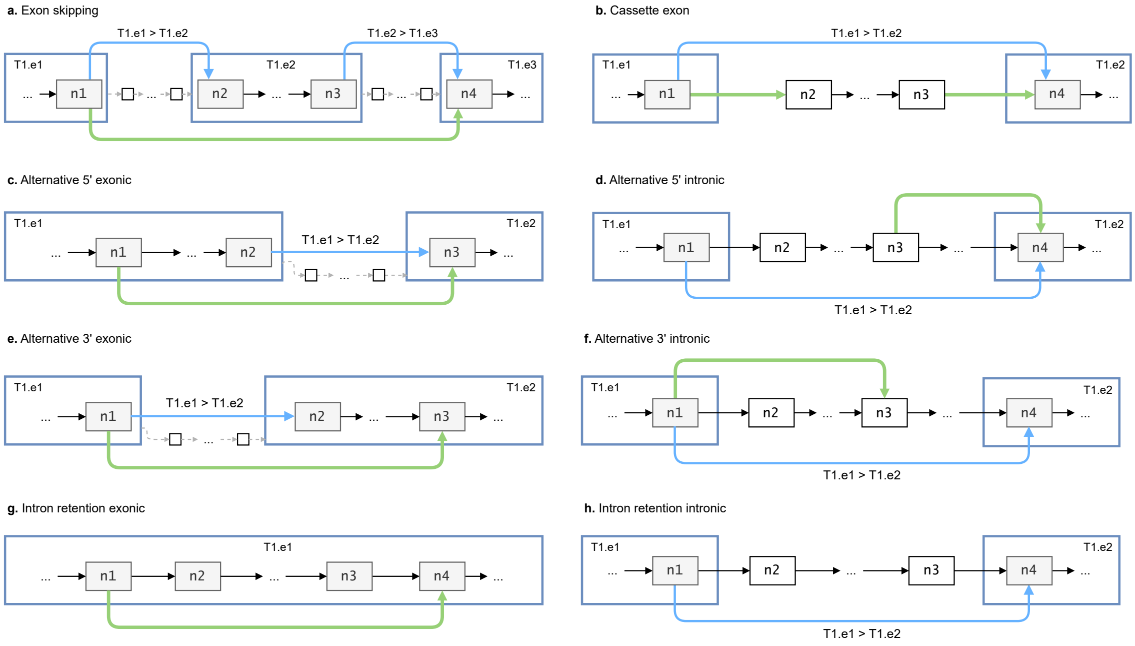


Fig. S1: All the novel events expressed within the annotated spliced pangenome showing the different tags used, weights are omitted for readability; blue squares represent exons with their tags, blue edges are the annotated junctions with their tags, green edges are the novel links, grey vertices are exonic and white are intronic,

| True Support ($\omega$) | Exon Skippings | Alternative 3' Sites | Alternative 5' Sites | Intron Retentions |
|---|---|---|---|---|
| 1 | 214 | 219 | 209 | 231 |
| 3 | 152 | 170 | 155 | 178 |
| 5 | 129 | 140 | 130 | 153 |
| 10 | 95 | 108 | 95 | 120 |
| 20 | 65 | 65 | 55 | 75 |

Table S1: Number of events reported by `asimulator` and supported by at least $\omega$ reads.

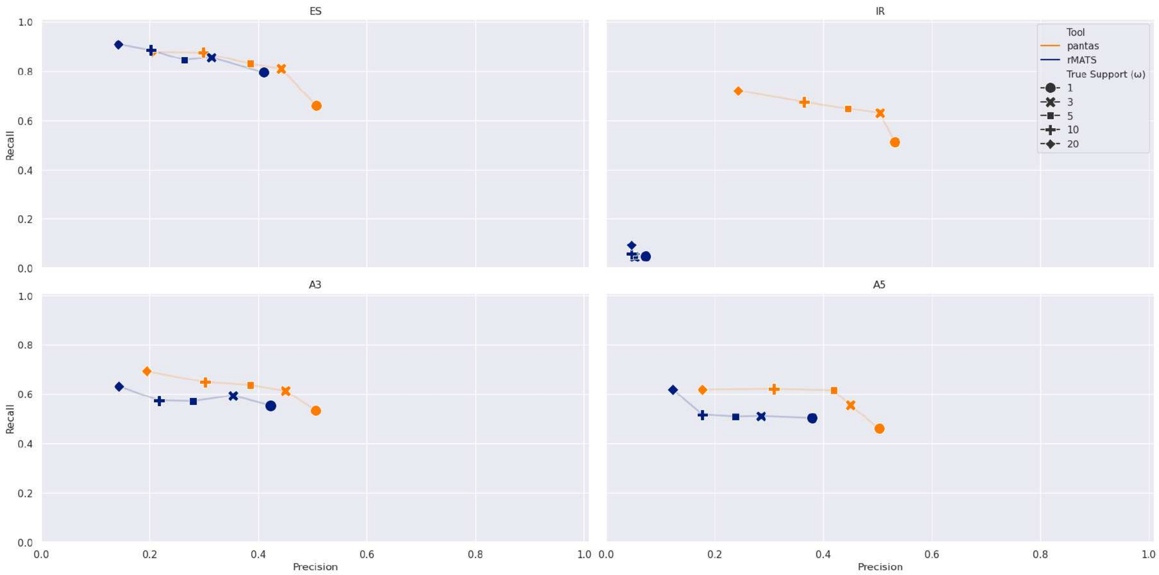


Fig. S2: Results on simulated data from Drosophila Melanogaster (novel events). Results are broken down by event type (ES: Exon Skipping, IR: Intron Retention, A3: Alternative 3', A5: Alternative 5').

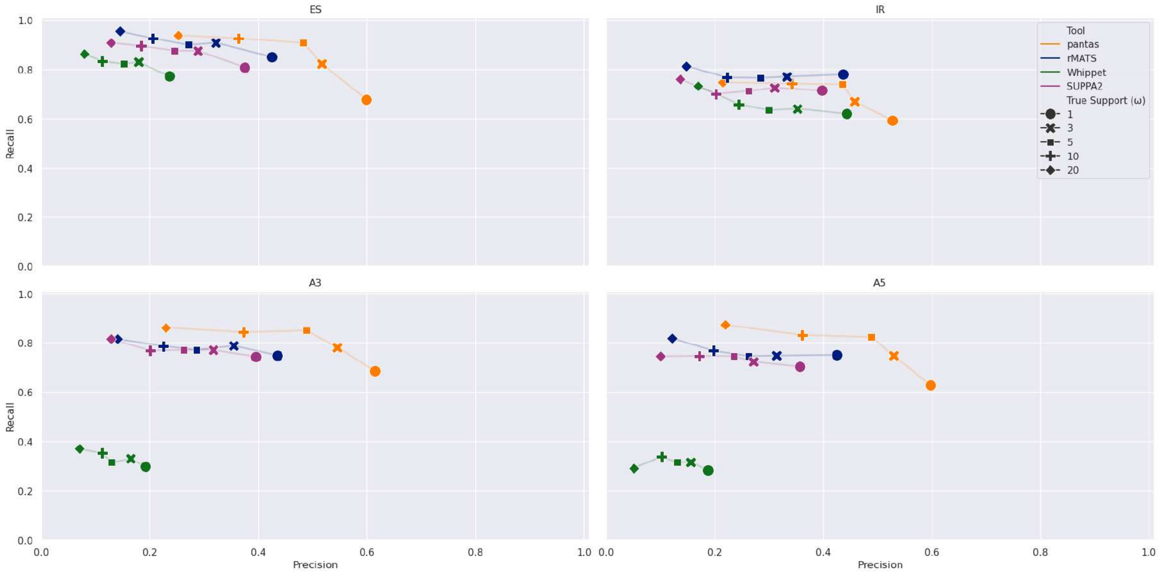


Fig. S3: Results on simulated data from Drosophila Melanogaster (annotated events). These results are obtained running `pantas` setting its $w$ parameter to 5 (minimum number of alignments needed to report an event). Results are broken down by event type (ES: Exon Skipping, IR: Intron Retention, A3: Alternative 3', A5: Alternative 5').

| ω | Event | Tool | TP | FN | FP | Prec | Rec | F1 |
|---|---|---|---|---|---|---|---|---|
| 1 | ES | pantas | 162 | 52 | 116 | 0.583 | 0.757 | 0.659 |
| 1 | ES | rMATS | 182 | 32 | 246 | 0.425 | 0.85 | 0.567 |
| 1 | ES | whippet | 165 | 49 | 534 | 0.236 | 0.771 | 0.361 |
| 1 | ES | SUPPA2 | 173 | 41 | 288 | 0.375 | 0.808 | 0.513 |
| 1 | IR | pantas | 150 | 81 | 141 | 0.515 | 0.649 | 0.575 |
| 1 | IR | rMATS | 180 | 51 | 232 | 0.437 | 0.779 | 0.56 |
| 1 | IR | whippet | 143 | 88 | 180 | 0.443 | 0.619 | 0.516 |
| 1 | IR | SUPPA2 | 165 | 66 | 250 | 0.398 | 0.714 | 0.511 |
| 1 | A3 | pantas | 164 | 55 | 104 | 0.612 | 0.749 | 0.674 |
| 1 | A3 | rMATS | 164 | 55 | 213 | 0.435 | 0.749 | 0.55 |
| 1 | A3 | whippet | 65 | 154 | 274 | 0.192 | 0.297 | 0.233 |
| 1 | A3 | SUPPA2 | 163 | 56 | 250 | 0.395 | 0.744 | 0.516 |
| 1 | A5 | pantas | 140 | 69 | 101 | 0.581 | 0.67 | 0.622 |
| 1 | A5 | rMATS | 157 | 52 | 212 | 0.425 | 0.751 | 0.543 |
| 1 | A5 | whippet | 59 | 150 | 255 | 0.188 | 0.282 | 0.226 |
| 1 | A5 | SUPPA2 | 147 | 62 | 265 | 0.357 | 0.703 | 0.473 |
| 3 | ES | pantas | 138 | 14 | 140 | 0.496 | 0.908 | 0.642 |
| 3 | ES | rMATS | 138 | 14 | 290 | 0.322 | 0.908 | 0.476 |
| 3 | ES | whippet | 126 | 26 | 573 | 0.18 | 0.829 | 0.296 |
| 3 | ES | SUPPA2 | 133 | 19 | 328 | 0.289 | 0.875 | 0.434 |
| 3 | IR | pantas | 129 | 49 | 162 | 0.443 | 0.725 | 0.55 |
| 3 | IR | rMATS | 137 | 41 | 275 | 0.333 | 0.77 | 0.464 |
| 3 | IR | whippet | 114 | 64 | 209 | 0.353 | 0.64 | 0.455 |
| 3 | IR | SUPPA2 | 129 | 49 | 286 | 0.311 | 0.725 | 0.435 |
| 3 | A3 | pantas | 145 | 25 | 123 | 0.541 | 0.853 | 0.662 |
| 3 | A3 | rMATS | 134 | 36 | 243 | 0.355 | 0.788 | 0.49 |
| 3 | A3 | whippet | 56 | 114 | 283 | 0.165 | 0.329 | 0.22 |
| 3 | A3 | SUPPA2 | 131 | 39 | 282 | 0.317 | 0.771 | 0.449 |
| 3 | A5 | pantas | 124 | 31 | 117 | 0.515 | 0.8 | 0.626 |
| 3 | A5 | rMATS | 116 | 39 | 253 | 0.314 | 0.748 | 0.443 |
| 3 | A5 | whippet | 49 | 106 | 265 | 0.156 | 0.316 | 0.209 |
| 3 | A5 | SUPPA2 | 112 | 43 | 300 | 0.272 | 0.723 | 0.395 |
| 5 | ES | pantas | 117 | 12 | 161 | 0.421 | 0.907 | 0.575 |
| 5 | ES | rMATS | 116 | 13 | 312 | 0.271 | 0.899 | 0.417 |
| 5 | ES | whippet | 106 | 23 | 593 | 0.152 | 0.822 | 0.256 |
| 5 | ES | SUPPA2 | 113 | 16 | 348 | 0.245 | 0.876 | 0.383 |
| 5 | IR | pantas | 110 | 43 | 181 | 0.378 | 0.719 | 0.495 |
| 5 | IR | rMATS | 117 | 36 | 295 | 0.284 | 0.765 | 0.414 |
| 5 | IR | whippet | 97 | 56 | 226 | 0.3 | 0.634 | 0.408 |
| 5 | IR | SUPPA2 | 109 | 44 | 306 | 0.263 | 0.712 | 0.384 |
| 5 | A3 | pantas | 119 | 21 | 149 | 0.444 | 0.85 | 0.583 |
| 5 | A3 | rMATS | 108 | 32 | 269 | 0.286 | 0.771 | 0.418 |
| 5 | A3 | whippet | 44 | 96 | 295 | 0.13 | 0.314 | 0.184 |
| 5 | A3 | SUPPA2 | 108 | 32 | 305 | 0.262 | 0.771 | 0.391 |
| 5 | A5 | pantas | 107 | 23 | 134 | 0.444 | 0.823 | 0.577 |
| 5 | A5 | rMATS | 97 | 33 | 272 | 0.263 | 0.746 | 0.389 |
| 5 | A5 | whippet | 41 | 89 | 273 | 0.131 | 0.315 | 0.185 |
| 5 | A5 | SUPPA2 | 97 | 33 | 315 | 0.235 | 0.746 | 0.358 |
| 10 | ES | pantas | 88 | 7 | 190 | 0.317 | 0.926 | 0.472 |
| 10 | ES | rMATS | 88 | 7 | 340 | 0.206 | 0.926 | 0.337 |
| 10 | ES | whippet | 79 | 16 | 620 | 0.113 | 0.832 | 0.199 |
| 10 | ES | SUPPA2 | 85 | 10 | 376 | 0.184 | 0.895 | 0.306 |
| 10 | IR | pantas | 87 | 33 | 204 | 0.299 | 0.725 | 0.423 |
| 10 | IR | rMATS | 92 | 28 | 320 | 0.223 | 0.767 | 0.346 |
| 10 | IR | whippet | 79 | 41 | 244 | 0.245 | 0.658 | 0.357 |
| 10 | IR | SUPPA2 | 84 | 36 | 331 | 0.202 | 0.7 | 0.314 |
| 10 | A3 | pantas | 91 | 17 | 177 | 0.34 | 0.843 | 0.484 |
| 10 | A3 | rMATS | 85 | 23 | 292 | 0.225 | 0.787 | 0.351 |
| 10 | A3 | whippet | 38 | 70 | 301 | 0.112 | 0.352 | 0.17 |
| 10 | A3 | SUPPA2 | 83 | 25 | 330 | 0.201 | 0.769 | 0.319 |
| 10 | A5 | pantas | 79 | 16 | 162 | 0.328 | 0.832 | 0.47 |
| 10 | A5 | rMATS | 73 | 22 | 296 | 0.198 | 0.768 | 0.315 |
| 10 | A5 | whippet | 32 | 63 | 282 | 0.102 | 0.337 | 0.156 |
| 10 | A5 | SUPPA2 | 71 | 24 | 341 | 0.172 | 0.747 | 0.28 |
| 20 | ES | pantas | 61 | 4 | 217 | 0.219 | 0.938 | 0.356 |
| 20 | ES | rMATS | 62 | 3 | 366 | 0.145 | 0.954 | 0.252 |
| 20 | ES | whippet | 56 | 9 | 643 | 0.08 | 0.862 | 0.147 |
| 20 | ES | SUPPA2 | 59 | 6 | 402 | 0.128 | 0.908 | 0.224 |
| 20 | IR | pantas | 57 | 18 | 234 | 0.196 | 0.76 | 0.311 |
| 20 | IR | rMATS | 61 | 14 | 351 | 0.148 | 0.813 | 0.251 |
| 20 | IR | whippet | 55 | 20 | 268 | 0.17 | 0.733 | 0.276 |
| 20 | IR | SUPPA2 | 57 | 18 | 358 | 0.137 | 0.76 | 0.233 |
| 20 | A3 | pantas | 56 | 9 | 212 | 0.209 | 0.862 | 0.336 |
| 20 | A3 | rMATS | 53 | 12 | 324 | 0.141 | 0.815 | 0.24 |
| 20 | A3 | whippet | 24 | 41 | 315 | 0.071 | 0.369 | 0.119 |
| 20 | A3 | SUPPA2 | 53 | 12 | 360 | 0.128 | 0.815 | 0.222 |
| 20 | A5 | pantas | 48 | 7 | 193 | 0.199 | 0.873 | 0.324 |
| 20 | A5 | rMATS | 45 | 10 | 324 | 0.122 | 0.818 | 0.212 |
| 20 | A5 | whippet | 16 | 39 | 298 | 0.051 | 0.291 | 0.087 |
| 20 | A5 | SUPPA2 | 41 | 14 | 371 | 0.1 | 0.745 | 0.176 |

Table S2: Full results on simulated data from Drosophila Melanogaster (annotated events). Precision, Recall, and F1 are reported for all true support threshold tested ($\omega$).

| $\omega$ | Event | Tool | TP | FN | FP | Prec | Rec | F1 |
|---|---|---|---|---|---|---|---|---|
| 1 | ES | pantas | 141 | 73 | 137 | 0.507 | 0.659 | 0.573 |
| 1 | ES | rMATS | 170 | 44 | 245 | 0.41 | 0.794 | 0.541 |
| 1 | IR | pantas | 118 | 113 | 104 | 0.532 | 0.511 | 0.521 |
| 1 | IR | rMATS | 11 | 220 | 139 | 0.073 | 0.048 | 0.058 |
| 1 | A3 | pantas | 117 | 102 | 114 | 0.506 | 0.534 | 0.52 |
| 1 | A3 | rMATS | 121 | 98 | 165 | 0.423 | 0.553 | 0.479 |
| 1 | A5 | pantas | 96 | 113 | 95 | 0.503 | 0.459 | 0.48 |
| 1 | A5 | rMATS | 105 | 104 | 171 | 0.38 | 0.502 | 0.433 |
| 3 | ES | pantas | 123 | 29 | 155 | 0.442 | 0.809 | 0.572 |
| 3 | ES | rMATS | 130 | 22 | 285 | 0.313 | 0.855 | 0.459 |
| 3 | IR | pantas | 112 | 66 | 110 | 0.505 | 0.629 | 0.56 |
| 3 | IR | rMATS | 8 | 170 | 142 | 0.053 | 0.045 | 0.049 |
| 3 | A3 | pantas | 104 | 66 | 127 | 0.45 | 0.612 | 0.519 |
| 3 | A3 | rMATS | 101 | 69 | 185 | 0.353 | 0.594 | 0.443 |
| 3 | A5 | pantas | 86 | 69 | 105 | 0.45 | 0.555 | 0.497 |
| 3 | A5 | rMATS | 79 | 76 | 197 | 0.286 | 0.51 | 0.367 |
| 5 | ES | pantas | 107 | 22 | 171 | 0.385 | 0.829 | 0.526 |
| 5 | ES | rMATS | 109 | 20 | 306 | 0.263 | 0.845 | 0.401 |
| 5 | IR | pantas | 99 | 54 | 123 | 0.446 | 0.647 | 0.528 |
| 5 | IR | rMATS | 8 | 145 | 142 | 0.053 | 0.052 | 0.053 |
| 5 | A3 | pantas | 89 | 51 | 142 | 0.385 | 0.636 | 0.48 |
| 5 | A3 | rMATS | 80 | 60 | 206 | 0.28 | 0.571 | 0.376 |
| 5 | A5 | pantas | 80 | 50 | 111 | 0.419 | 0.615 | 0.498 |
| 5 | A5 | rMATS | 66 | 64 | 210 | 0.239 | 0.508 | 0.325 |
| 10 | ES | pantas | 83 | 12 | 195 | 0.299 | 0.874 | 0.445 |
| 10 | ES | rMATS | 84 | 11 | 331 | 0.202 | 0.884 | 0.329 |
| 10 | IR | pantas | 81 | 39 | 141 | 0.365 | 0.675 | 0.474 |
| 10 | IR | rMATS | 7 | 113 | 143 | 0.047 | 0.058 | 0.052 |
| 10 | A3 | pantas | 70 | 38 | 162 | 0.302 | 0.648 | 0.412 |
| 10 | A3 | rMATS | 62 | 46 | 224 | 0.217 | 0.574 | 0.315 |
| 10 | A5 | pantas | 59 | 36 | 132 | 0.309 | 0.621 | 0.413 |
| 10 | A5 | rMATS | 49 | 46 | 227 | 0.178 | 0.516 | 0.264 |
| 20 | ES | pantas | 57 | 8 | 221 | 0.205 | 0.877 | 0.332 |
| 20 | ES | rMATS | 59 | 6 | 356 | 0.142 | 0.908 | 0.246 |
| 20 | IR | pantas | 54 | 21 | 168 | 0.243 | 0.72 | 0.364 |
| 20 | IR | rMATS | 7 | 68 | 143 | 0.047 | 0.093 | 0.062 |
| 20 | A3 | pantas | 45 | 20 | 187 | 0.194 | 0.692 | 0.303 |
| 20 | A3 | rMATS | 41 | 24 | 245 | 0.143 | 0.631 | 0.234 |
| 20 | A5 | pantas | 34 | 21 | 158 | 0.177 | 0.618 | 0.275 |
| 20 | A5 | rMATS | 34 | 21 | 242 | 0.123 | 0.618 | 0.205 |

Table S3: Full results on simulated data from Drosophila Melanogaster (novel events). Precision, Recall, and F1 are reported for all true support threshold tested ($\omega$).

| Event | pantas | rMATS | whippet | SUPPA2 |
|---|---|---|---|---|
| *ES* | 370 | 366 | 3205 | 140 |
| *A3* | 337 | 200 | 401 | 129 |
| *A5* | 367 | 189 | 385 | 131 |
| *IR* | 322 | 177 | 331 | 113 |
| ***All*** | 1396 | 932 | 4322 | 513 |

Table S4: Alternative splicing events count for Drosophila Melanogaster experiments on real data.

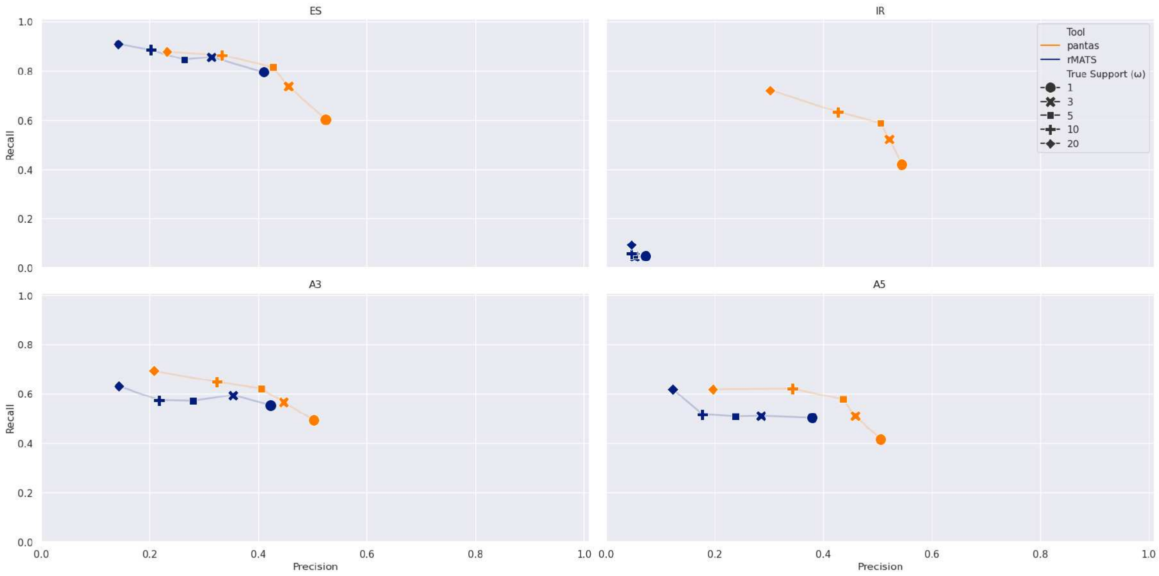


Fig. S4: Results on simulated data from Drosophila Melanogaster (novel events). These results are obtained running `pantas` setting its $w$ parameter to 5 (minimum number of alignments needed to report an event). Results are broken down by event type (ES: Exon Skipping, IR: Intron Retention, A3: Alternative 3', A5: Alternative 5').

| **Event** | `pantas` | `rMATS` | `whippet` | `SUPPA2` |
|---|---|---|---|---|
| *ES* | 553 | 1220 | 12030 | 755 |
| *A3* | 627 | 754 | 731 | 944 |
| *A5* | 630 | 638 | 705 | 909 |
| *IR* | 506 | 536 | 537 | 663 |
| ***All*** | 2316 | 3148 | 14003 | 3271 |

Table S5: AS events count for Drosophila Melanogaster experiments without any filtering on *minimum* $\Delta\psi$, *minimum probability*, and *p-value*.

| Tool | Time (min.) | RAM (GB) |
|---|---|---|
| vg index | 128 | 20 |
| vg mpmap | 388* | 3 |
| pantas weight | 26* | 5 |
| pantas call | 11* | 16 |
| pantas quant | 1 | 1 |
| STAR index | 1 | 4 |
| STAR align | 20* | 6 |
| rMATS | 2 | 1 |
| salmon index | 0.5 | 1 |
| salmon quant | 1* | 1 |
| SUPPA2 | 0.5 | 1 |
| whippet index | 2 | 1 |
| whippet quant | 9* | 1 |
| whippet delta | 1 | 1 |

* average over the 6 samples.

Table S6: Efficiency results on real data from Drosophila Melanogaster. Time usage is reported in minutes. Peak RAM usage is reported in GB.

| Tool | Time (min) | RAM (GB) |
|---|---|---|
| `shark` | 2* | 2 |
| `vg index` | 2 | 8 |
| `vg mpmap` | 20* | 3 |
| `pantas weight` | 1* | 1 |
| `pantas call` | 1* | 1 |
| `pantas quant` | 1 | 1 |
| STAR index | 43 | 38 |
| STAR align | 9* | 32 |
| rMATS | 9 | 2 |
| salmon index | 2 | 1 |
| salmon quant | 2* | 3 |
| SUPPA2 | 13 | 1 |
| whippet index | 45 | 5 |
| whippet quant | 29* | 6 |
| whippet delta | 2 | 1 |

* average over the 6 samples.

Table S7: Efficiency results on real Human data. Time usage is reported in minutes. Peak RAM usage is reported in GB.

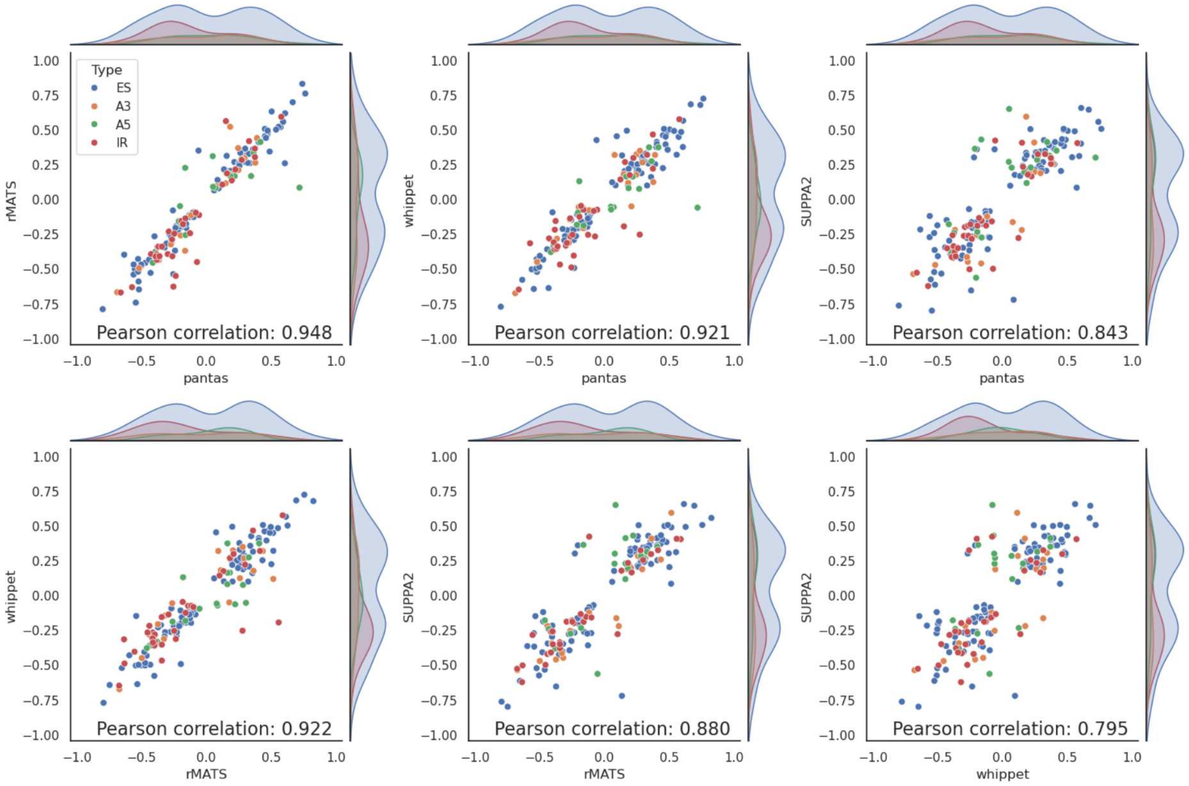


Fig. S5: Results on real data from Drosophila Melanogaster. All-vs-all correlation plots of the $\Delta\psi$ reported by the considered tools for the 164 events shared among the 4 tools. Each point is an event color coded based on its type.

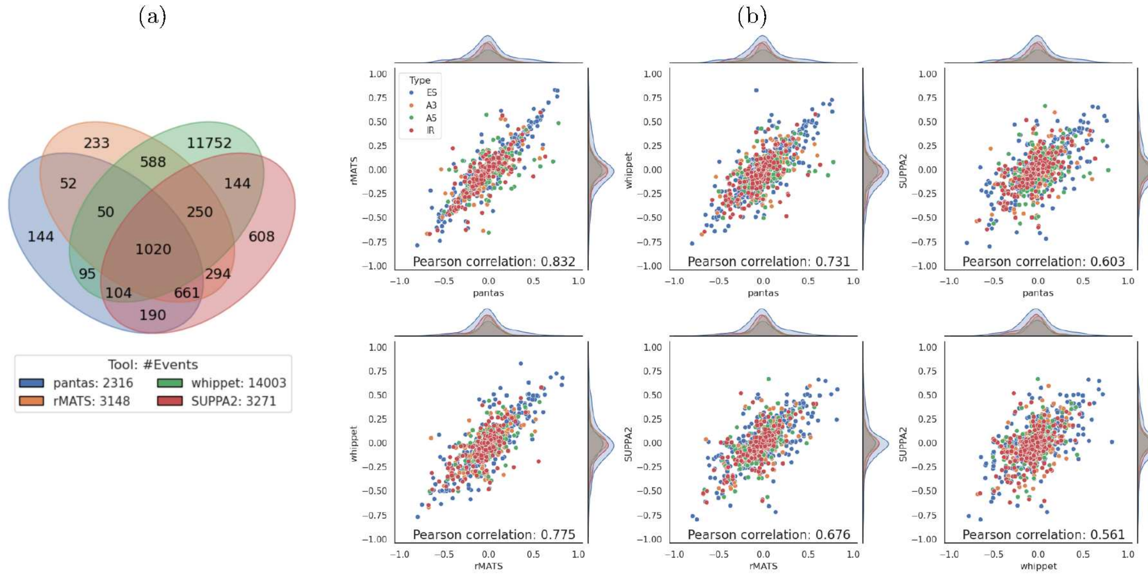


Fig. S6: Results on real data from Drosophila Melanogaster (all events). (a) Venn diagram showing the number of AS events reported by each tool. (b) All-vs-all correlation plots of the $\Delta\psi$ reported by the considered tools for the 1020 events shared among the 4 tools. Each point is an event color coded based on its type.

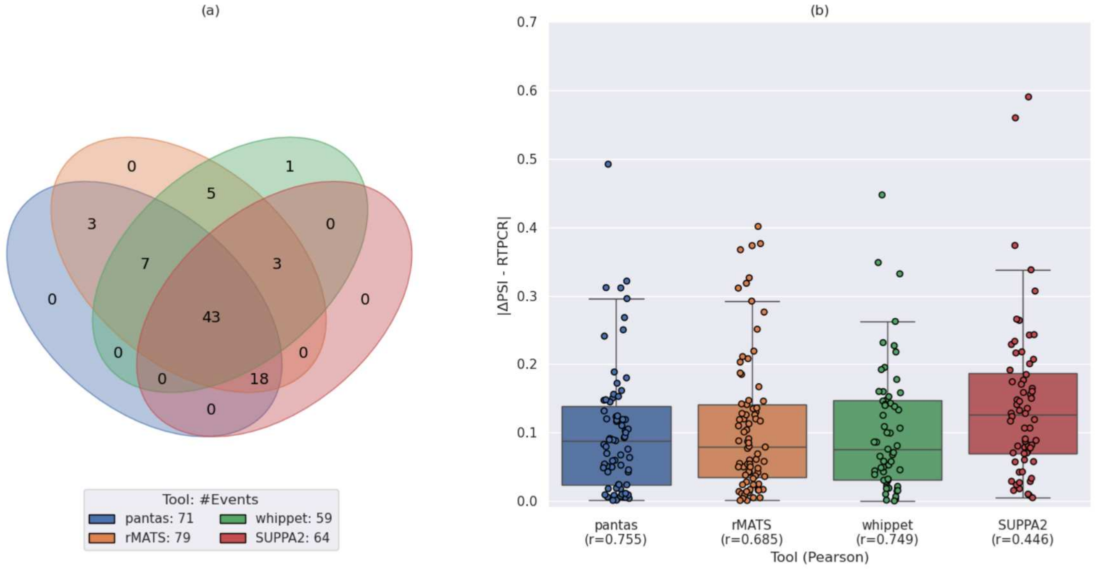


Fig. S7: (a) Venn diagram showing the number of AS events reported by each tool. The legend reports the total number of events reported by the tool. (b) Boxplot showing the distribution of the difference between the $\Delta\psi$ predicted by each tool and the $\Delta\psi$ provided by RT-PCR. The x axis labels report the tool name and the Pearson correlation ($r$) between the predicted $\Delta\psi$ and the RT-PCR $\Delta\psi$.

.
